# Integrator Subunit 4 is a ‘Symplekin-like’ Scaffold that Associates with INTS9/11 to Form the Integrator Cleavage Module

**DOI:** 10.1101/172635

**Authors:** Todd R. Albrecht, Sergey P. Shevtsov, Lauren G. Mascibroda, Natoya J. Peart, Iain A. Sawyer, Miroslav Dundr, Eric J. Wagner

**Affiliations:** Department of Biochemistry and Molecular Biology, University of Texas Medical Branch at Galveston, Galveston TX 77550; Department of Cell Biology, Rosalind Franklin University of Medicine and Science, Chicago Medical School, North Chicago IL 60064; Laboratory of Receptor Biology and Gene Expression, National Cancer Institute, National Institutes of Health, Bethesda MD 20892

**Keywords:** Integrator Complex, snRNA, Symplekin, histone mRNA processing, Cajal bodies, Histone locus bodies

## Abstract

Integrator (INT) is a transcriptional regulatory complex associated with RNA polymerase II that is required for the 3’-end processing of both UsnRNAs and enhancer RNAs. Integrator subunits 9 (INTS9) and INTS11 constitute the catalytic core of INT and are paralogues of the cleavage and polyadenylation specificity factors CPSF100 and CPSF73. While CPSF73/100 are known to associate with a third protein called Symplekin, there is no paralog of Symplekin within INT raising the question of how INTS9/11 associate with the other INT subunits. Here, we have identified that INTS4 is a specific and conserved interaction partner of INTS9/11 that does not interact with either subunit individually. Although INTS4 has no significant homology with Symplekin, it possesses N-terminal HEAT repeats similar to Symplekin but also contains a β-sheet rich C-terminal region, both of which are important to bind INTS9/11. We assess three functions of INT including UsnRNA 3’-end processing, maintenance of Cajal body integrity, and formation of histone locus bodies to conclude that INTS4/9/11 are the most critical of the INT subunits for UsnRNA biogenesis. Altogether, these results indicate that INTS4/9/11 compose a heterotrimeric complex that likely represents the Integrator ‘cleavage module’ responsible for its endonucleolytic activity.

## Introduction

The Integrator Complex (INT) is a metazoan-specific group of proteins comprised of at least 14 subunits that associates with the phosphorylated C-terminal domain (CTD) of the largest subunit of RNA polymerase II (RNAPII)(reviewed in (1)). Through its association with RNAPII, one of the functions of INT is to cleave the extended 3’-end of Uridine-rich small nuclear RNAs (UsnRNAs) and this process is essential for the biogenesis of spliceosomal snRNPs (2,3). Recently, INT has also been shown to be involved in the 3’-end formation of enhancer RNA (eRNA) thereby impacting the transcriptional activation of neighboring protein-encoding genes (4). In addition to these functions as a 3’-end RNA processing factor, ChIP-seq analyses of INT in human cells indicate that INT is associated with the promoters of a large number of protein-encoding genes and is significantly enriched in genes that harbor paused RNAPII (5-7). The proposed purpose of INT occupancy at these promoters is to promote pause-release and, indeed it has been demonstrated that epidermal growth factor (EGF)-responsive immediate early genes in mammalian cells and heat shock response genes in *Drosophila* both require INT for productive release of RNAPII. These broadly important functions in gene regulation are reflected by a diverse collection of cellular processes that are perturbed under conditions of INT dysfunction. Several INT subunits have been found to play important roles in adipose tissue differentiation, embryonic development, ciliogenesis, lung homeostasis, structural integrity of a major nuclear feature, the Cajal body (CB), as well as in the formation of herpesvirus microRNA 3’-end formation (8-16). More recently, mutations of Integrator subunits 1 (INTS1) and INTS8 have been implicated in disrupted human brain development and a large-scale analysis of TCGA database data reveals significant misexpression of INT subunits in a spectrum of human tumor types (17,18).

Despite its importance to gene expression, we have little understanding of how INT is constructed and the function of the majority of the individual subunits has not been determined. Our current understanding of INT function and assembly is based, in part, on deriving information from related subunits of the cleavage and polyadenylation machinery (reviewed in (19)). For the processing of either pre-mRNA or histone pre-mRNA, endonucleolytic cleavage is carried out by the cleavage and polyadenylation specificity factor of 73kDa (CPSF73)(20,21), which is a member of the metallo-β-lactamase/ β-CASP family of RNA endonucleases (22). CPSF73 is known to associate with its heterodimeric partner, CPSF100 and, together with a third protein called Symplekin form what has been termed a Core Cleavage Complex (CCC)(23,24). In the case of eukaryotic pre-mRNA processing, these three subunits then interact with other members of the processing complex to catalyze cleavage of the pre-mRNA. Specifically, Symplekin has been shown to interact with several other subunits of the processing machinery including CstF64 and Ssu72 with the latter ultimately associating with the C-terminal domain of Rpb1 (25-29). The *Saccharomyces cerevisae* homolog of Symplekin, Pta1, has also been shown to perform a similar function of a scaffolding subunit and has been found to interact both with members of the CF and CPF complex (30). Not surprisingly, depletion of Symplekin from *Drosophila* S2 cells leads to profound misprocessing of histone premRNA and the degree of the misprocessing is similar to that which is observed upon depletion of CPSF73 and CPSF100 (24,31).

The high degree of conservation between CPSF73 and CPSF100 with INTS11 and INTS9 respectively, was instrumental in providing clues that INT is also an RNA endonuclease (2). INTS11 is 40% identical to CPSF73 throughout their β-lactamase and β-CASP domains while both CPSF100 and INTS9 contain alterations in critical zinc-coordinating residues rendering the two enzymes to be predicted inactive (32). Similar to its CPSF counterparts, INTS9 and INTS11 utilize their C-terminal domains (CTD) to form a heterodimer and this is critical to their role in UsnRNA biogenesis (33). The important difference between the CPSF and INT heterodimer is that their CTDs are highly divergent. We recently solved the crystal structure of the CTDs of both INTS9 and INTS11 in a complex and found that the CTDs of both subunits form a common domain created upon heterodimerization and this structure is essential for INT function (34). One prediction that this observation generates is that another INT subunit recognizes the heterodimeric INTS9/11 domain to either stabilize it and/or allow its association with the other INT subunits. The potential implication of this model is that the CTDs of CPSF73/100 also form a common domain upon heterodimerization that is topologically distinct from INTS9/11 and recognized by Symplekin. Binding under the requirement of heterodimerization would ensure that only *bona fide* cleavage factors are incorporated into each of the processing complexes. Unfortunately, beyond INTS9 and INTS11 there are no other members of INT that resemble members of the cleavage and polyadenylation machinery and no candidate ‘Symplekin-like’ INT subunit can be readily identified using homology modeling. This suggests that either INTS9/11 associates with other members of INT using a mechanism unrelated to how Symplekin functions in pre-mRNA processing or that the Symplekin-like subunit within INT has diverged enough to prevent homology-based prediction.

Many RNA species processed by INT are accumulated in specialized nuclear structures known as nuclear bodies. Indeed, the close relationship between specific gene transcription, RNA processing and nuclear body function is well-established (35-39) and several groups have shown that artificially-induced RNA accumulation, representative of gene expression, is capable of nucleating several nuclear bodies *de novo* (40,41). Additionally, depletion of the CB, which functions as a general INT-dependent RNP assembly and modification factory, decreases the expression of all classes of INT-processed UsnRNAs and INT-independent small nucleolar RNAs (snoRNAs) and CB-specific RNAs (scaRNAs)(42). This is likely due to the enrichment of various transcriptional and assembly regulators within the CB and other, to be determined, factors (43). Interestingly, the CB has been shown to physically associate with another nuclear body, the histone locus body (HLB), which may accelerate the delivery of the INT-processed low abundant U7 snRNA, which is matured in the CB and catalyzes the 3’-end trimming of histone pre-mRNAs in the HLB (42,44). However, despite limited evidence linking some INT subunit depletion to CB disassembly (15), no information exists for the remaining INT subunit-dependency of the CB, endogenous INT localization relative to the CB, nor the downstream effects on HLB stability.

Here, we have developed a modified yeast two-hybrid screen to determine that INTS4 binds to the INTS9/11 heterodimer and that this binding is conserved from *Drosophila* to humans. INTS4 binding to INTS9/11 is intimately dependent upon their heterodimerization and both N-terminal and C-terminal regions of INTS4 are involved in the interaction. We demonstrate that depletion of any of these three subunits causes the most robust amount of UsnRNA misprocessing using a quantitative reporter system indicating that these three subunits not only physically interact but also provide an equivalent functional requirement that exceeds other INT subunits tested. Consistently, depletion of INTS4/9/11 by siRNA in HeLa cells also causes complete disassembly of both Cajal bodies and, subsequently, histone locus bodies, two major and frequently physically associated nuclear bodies which are primarily nucleated around snRNA and clustered replication-dependent histone genes and are involved in snRNA biogenesis and histone mRNA production. These data strongly support the model that these three subunits comprise a minimal Integrator cleavage module and ongoing spliceosomal snRNP biogenesis through the activity of INTS4/9/11 is critical to maintain these nuclear structures.

## Materials and Methods

### Modified yeast two-hybrid analysis

Yeast two-hybrid assays were carried out in PJ69-4a and PJ49-4α strains. Human INTS11 or INTS9 CTD fragments were cloned into either pOBD, pOAD or pTEF vectors using conventional cloning. Clones were sequenced to verify identity; PCR primers are available upon request. pOBD plasmids were transformed into PJ69-4a yeast and were selected on tryptophan-dropout medium; pOAD plasmids were transformed into PJ49-4α yeast and were selected on leucine-dropout medium. pTEF vectors were transformed in PJ69-4a yeast containing specific pOBD vectors and were selected on tryptophan, uracil-dropout medium. Mating the yeast strains followed by selection on medium lacking tryptophan, uracil and leucine created triple transformants. Interactions were tested through serial dilution of diploid yeast followed by plating on medium lacking tryptophan, uracil and leucine or on medium lacking tryptophan, uracil, leucine, and histidine that also was supplemented with 5 mM 3-amino-1,2,4-triazole.

### Cell culture and reporter transfections

HeLa human cervical carcinoma cells were cultured using Dulbecco’s modified Eagle medium (DMEM) (Invitrogen) supplemented with 10% fetal bovine serum (FBS; Phenix Research) and penicillin-streptomycin (pen/strep) (Invitrogen). Integrator knockdown transfections were performed using HeLa cells using either RNAiMAX (Invitrogen) or Lipofectamine 2000 (Invitrogen) following previously described protocols (45). Briefly, 1 X 10^5^ HeLa cells were seeded in a 24-well plate (Greiner Bio-One, Germany) 1 day prior to transfection. Day 1, 10µm of siRNA was mixed in 50 µl of Opti-MEM (Invitrogen) without serum. A second tube with 50 µl of Opti-MEM was mixed with 2 µl of RNAiMAX and allowed to incubate at room temperature for 7 min. The contents of the two tubes were mixed and allowed to incubate for 25 min at room temperature. Day 2, cells were trypsinized and plated in a 6-well plate (Greiner Bio-One, Germany). Day 3, 10µm of siRNA was mixed in 50 µl of Opti-MEM (Invitrogen) without serum. A second tube with 50 µl of Opti-MEM was mixed with 2 µl of RNAiMAX and allowed to incubate at room temperature for 7 min. The contents of the two tubes were mixed and allowed to incubate for 25 min at room temperature. Day 4, 100 µg of U7-GFP reporter and 500µg of INT plasmids was mixed in 100 µl of Opti-MEM (Invitrogen) without serum. A second tube with 100 µl of Opti-MEM was mixed with 2 µl of Lipofectamine 2000 and allowed to incubate at room temperature for 7 min. The contents of the two tubes were mixed and allowed to incubate for 25 min at room temperature. The entire 200 µl mixture was directly added to the cells that had been plated in 1 ml of growth media containing 10% FBS. Media were changed 24 h later depending on cell fitness and ultimately harvested 48 h after the initial transfection. The sequences for the siRNA used in this study are as follows: INTS1 (#1-5’ 3’, #2-5’CAUUUCUCCGUCGAUUAAA3’), INTS3 (#1-5’GAAGUACUGAGUUCAGAUA3’, #2-5’CAGAAAGUGUUCUGGAUAU3’), INTS4 (#1-5’GUAGGCUUAAGGAGUAUGUGA3’, #2-5’GUAGGCUUAAGGAGUAUGUGA3’), INTS7 (#1-5’GGCTAAATAGTTTGAAGGA3’, #2-5’ 3’), INTS9 (#1-5’GAAATGCTTTCTTGGACAA3’, #2-5’ 3’), INTS10 (#1-5’ GGATACTTGGCTTTGGTTA3’, #2-5’ 3’), INTS11 (#1-5’GACAGACCCUGGGCCUGCA3’, #2-5’ CAGACUUCCUGGACUGUGU3’), INTS12 (#1-5’GTCAAGACATCCACAGTTA3’, #2-5’ 3’). Control and SLBP siRNA were described previously (46).

### Immunoblot analysis

Western blots were performed using SDS/PAGE as described previously (33). Western blots were conducted using antibodies raised to INTS1 (Bethyl), INTS3 (Bethyl), INTS4 (Bethyl), INTS7 (Bethyl), INTS9 (Bethyl), INTS10 (Bethyl), INTS11 (Bethyl), INTS12 (Bethyl), tubulin (Abcam), Coilin (Santa Cruz), SMN (Santa Cr), Gemin2 (Santa Cruz), GFP (Clontech) and FLAG epitope (Sigma).

### Immunofluorescence and confocal microscopy

HeLa cells were grown on glass coverslips in Dulbecco’s modified eagle’s medium (DMEM) (Invitrogen) supplemented with 10% (v/v) fetal calf serum (FCS; Invitrogen) and 1% glutamine, penicillin and streptomycin, and 5% CO_2_ at 37°C. Cells were washed with PBS and fixed with 2% paraformaldehyde in PBS for 10 min at room temperature, then washed with PBS and permeabilized with 0.2% Triton X-100 in PBS on ice, and again washed with PBS. Cells were immunolabelled using specific antibodies against INT subunits (1:200), coilin (1:400, H-300, Santa Cruz, or 1:200, pDelta, Sigma), SMN (1:200, Santa Cruz), Gemin2 (1:200, Santa Cruz), NPAT (provided by Dr. J. Zhao, University of Rochester Medical Center, USA), Fibrillarin (1:100, 72B9, provided by Dr. J. Gall, Carnegie Institution for Science, USA), SLBP (1:100, 2C4, Sigma). Primary antibodies were detected with appropriate secondary antibodies conjugated with FITC or Cy5 (Jackson Laboratories). Cells were mounted using ProLong Gold (Invitrogen) and observed on Zeiss LSM510 or LSM780 confocal microscopes using 63x 1.4 N.A. or 100x 1.4 N.A. Plan Apochromat oil objectives. Images were collected as vertical *z*-stacks (*z*; 200 nm) and projected as maximum intensity projections.

## Results

### A modified yeast two-hybrid screen identifies INTS4 as an interactor of the INS9/INTS11 heterodimer

We previously established that heterodimeric interaction between INTS9 and INTS11 could be detected and measured using yeast two-hybrid analysis (34) and that this interaction is highly robust with resistance to 3-amino-1,2,4-triazole (3-AT) supplementation to 100mM concentration (47). Given this interaction and our detailed understanding of the structural basis for heterodimerization, we designed a method to create a strain of yeast where the INTS9/11 heterodimer could be used as a bait to identify other potential INT subunits that bind to these proteins once associated (Fig. 1A). To that end, we fused human INTS9 protein to the Gal4 DNA binding domain, which is selected for in yeast using a tryptophan marker and then expressed full-length human INTS11 as an unfused protein *in trans* using a uracil marker. In this strain, we then individually expressed the other 14 INT subunits fused to the Gal4 activation domain and grew the yeast either on permissive media or on selective media that lacked histidine and was supplemented with 3-AT to test for potential interactions. We observed that none of the INT subunits could support growth on selective medium with the exception of INTS4 (Fig. 1A). The growth observed on selective media is observed either because INTS4 binds to the INTS9/11 heterodimer or because INTS4 binds only to INTS9 irrespective of INTS11. To address this, we conducted a second screen where INTS11 was fused to the DNA binding domain and INTS9 is expressed as an unfused protein *in trans*. Using this configuration, we again observed that only INTS4 could support growth on selective medium (Fig. 1B). Finally, we repeated both sets of screens using a collection of cDNAs derived entirely from *Drosophila* where all fourteen INT subunits have been previously identified functionally and since annotated (3,48). In complete agreement with the mammalian screens, we observe that only *Drosophila* INTS4 is capable of binding to the dINTS9/11 heterodimer regardless of the position of the two subunits in the screen configuration (Figs. 1C/D).

**Figure 1.**
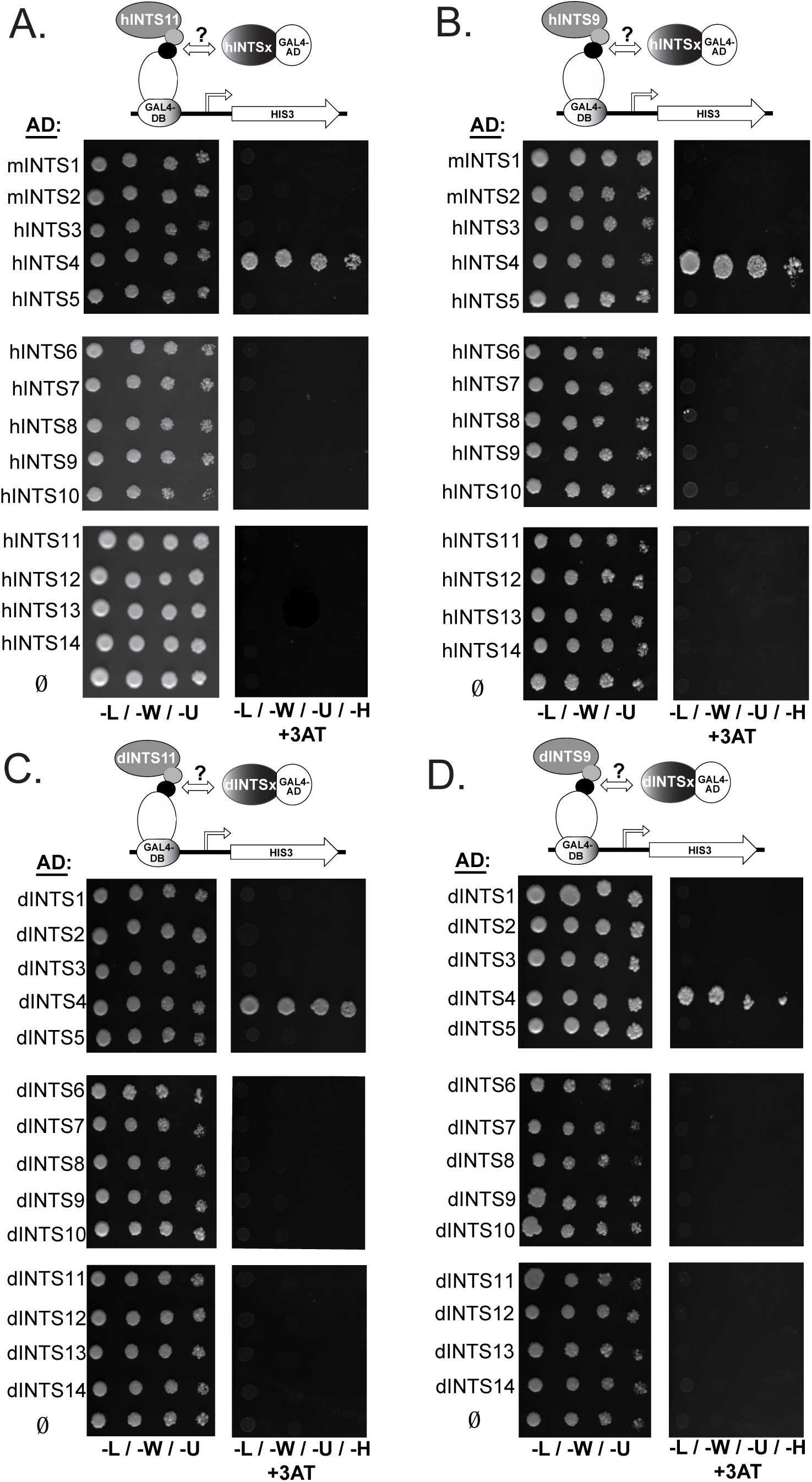
Multiple modified yeast two-hybrid screens identify INTS4 as an INTS9/11 interacting protein. A. (Upper panel) Human INTS9 is fused to the Gal4 DNA binding domain (DB) and is co-expressed with human INTS11 that is not fused to any yeast protein and was screened against an activation domain library containing each of the mammalian INT subunits. (Lower panel) Results of the modified yeast two-hybrid screen identifying INTS4 as the only INT subunit capable of supporting growth on selection media lacking histidine and supplemented with 3-AT. B (Upper panel) Human INTS11 is fused to the Gal4 DNA binding domain and co-expressed with human INTS9 that is not fused to any yeast protein and was screened against an activation domain library containing each of the mammalian INT subunits. C/D. Same as panels A and B except both screens were conducted using INT subunits derived from Drosophila melanogaster.

To more rigorously address the hypothesis that INTS4 binds to the INTS9/11 heterodimer, we proceeded to test the interaction using a series of additional controls. We first repeated the screen, but in each case, we removed one of the three components from the different positions to rule against potential auto-activation. The results of this approach demonstrate that replacement of any of the three INT-encoding plasmids with empty vectors resulted in loss of interaction demonstrating that auto-activation is not occurring and that all three subunits are required for growth (Fig. 2A). These same controls were introduced into plasmids encoding *Drosophila* INTS4/9/11 and near identical results were attained with the exception that the combination of subunits where dINTS11 is fused to the Gal4 DNA binding domain reproducibly resulted in the lowest (but clearly detectable) amount of growth of yeast on selective medium (Fig. 2B). A second set of control constructs stems from our recent crystal structure of the INTS9/11 heterodimer where we identified single or double amino acid mutations capable of disrupting their interaction (34). Importantly, these mutations, when introduced into the full-length proteins, are capable of abrogating the association of INTS9/11 using coimmunoprecipitation assays but have no effect on protein stability or accumulation. With this information in hand, we mutated human INTS9 by changing arginine-644 to a glutamate in the context of INTS9 fused to the Gal4 DNA binding domain and INTS11 expressed *in trans*. We observed that the INTS9 mutation completely eliminated the ability of INTS4 to support growth on selective medium (Fig. 2C). Conversely, when two mutations were introduced into INTS11 (L509A/F511A) in the context of INTS11 expressed as a DNA binding fusion, we also observed a total loss of growth on selective medium (Fig. 2D). Altogether, these results are highly suggestive that INTS4 binds to INTS9 and INTS11 only when they are associated with each other through heterodimerization of their C-terminal domains.

**Figure 2.**
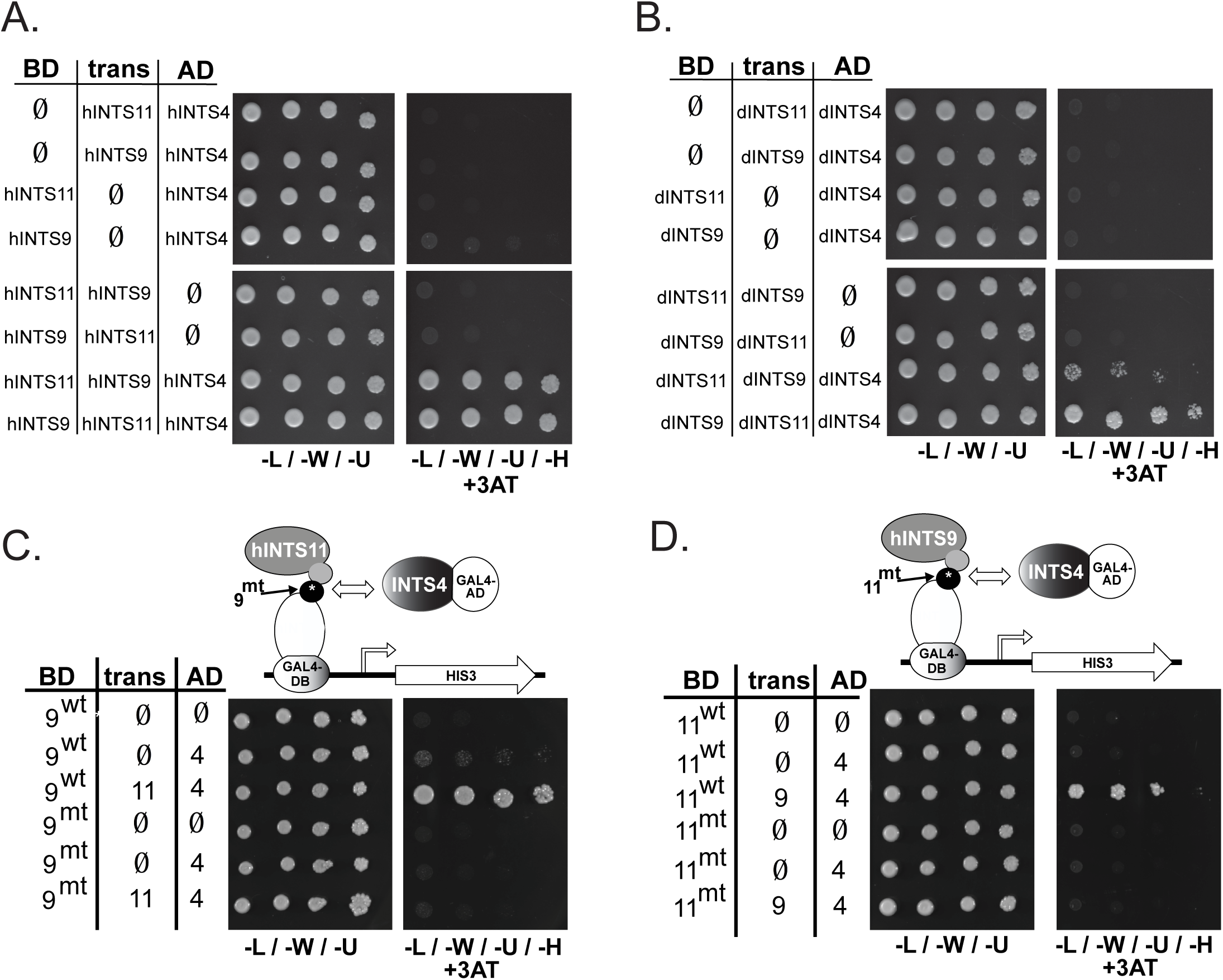
INTS4 likely recognizes the INTS9/11 heterodimeric interface to mediate interaction. A. Results of modified yeast two-hybrid where one of the subunits is eliminated at each position. “BD” represents the INT subunit fused to the DNA binding domain; “trans” represents the INT subunit expressed as a non-fused protein; “AD” represents the INT subunit expressed as an activation domain fusion protein. B. Same as panel A except using Drosophila components. C. Results of modified yeast two-hybrid using INTS9 as the DNA binding fusion protein but with the introduction of a mutation (mt) that disrupts the INTS9/11 heterodimer. D. Same as in panel D except the position of INTS11 and INTS9 are switched and the mutation is introduced into INTS11.

### The INTS4 N- and C-terminal regions are required for INTS9/11 interaction and INTS4 function

While there is no significant homology detected between Symplekin and INTS4, there are some notable similarities and differences in domains present in the two proteins. The structure of the Symplekin N-terminus is known (49) and has been shown to possess seven HEAT repeats that interact with Ssu72 (29), which in turn binds to the CTD of RNAPII. The region of Symplekin that binds to CPSF73/100 has not been as well characterized but includes a large stretch of the C-terminus, from amino acids 272-1080 of *Drosophila* Symplekin (23). In the case of INTS4, structure prediction programs identified an array of N-terminal HEAT repeats that may number as many as ten over the first 800 amino acids while the remaining 200 amino acids at the C-terminal are predicted to be rich in β-sheets (Fig. 3A). Using this information as a guide, we tested what regions of INTS4 are required for its interaction with INTS9/11. We first deleted N-terminal HEAT repeats, two at a time and observed that removal of just the first two HEAT repeats resulted in a reduction in the growth of yeast on selective medium (Fig. 3B). Further deletions from the first four HEAT repeats to removal of all ten resulted in no growth (Fig. 3B). These results indicate that the first two HEAT repeats are likely required for full binding to INTS9/11 and that the first four HEAT repeats are absolutely essential to mediate interaction. Additionally, we deleted regions of INTS4 starting from the C-terminus and found that removal of the last 100 amino acids also resulted in the loss of growth on selective medium (Fig. 3C) demonstrating that the N-terminal HEAT repeats are not sufficient to mediate interaction and that more than one region of INTS4 associates with INTS9/11. To attain a higher resolution analysis of the C-terminus of INTS4, we designed smaller deletions of only 5 amino acids at a time. Using this approach, we found that removal of only the last ten amino acids of INTS4 in the context of the full-length protein was sufficient to disrupt growth on selectable media (Fig. 3D). Finally, we conducted an alanine scanning analysis of the last 40 amino acids of INTS4 where five amino acids at a time were changed to alanine to generate eight total mutants. Using this approach, we observed that three distinct but consecutive sets of alanine mutants ranging from amino acids 944-958 are required for interaction with INTS9/11 while all other mutants that were constructed failed to impact growth on selective medium (Fig. 3E). These results indicate that sequences within both the N-terminus and C-terminus of INTS4 are required to interact with INTS9/11.

**Figure 3.**
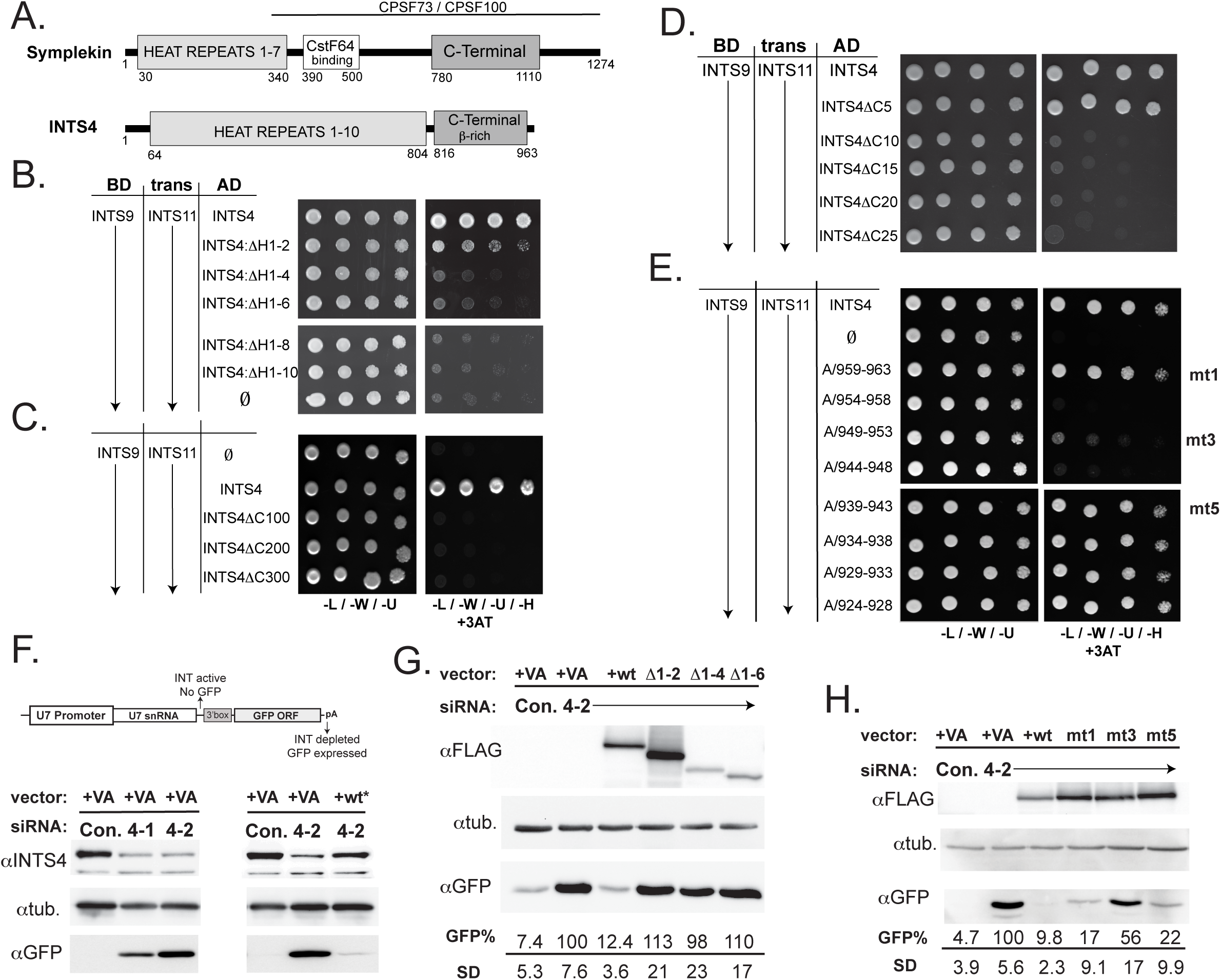
Regions within the N- and C-terminus are required for INTS4 binding and are also essential for INTS4 function in UsnRNA processing. A. Schematic of human Symplekin and INTS4 highlighting known and predicted domains. B. Results of modified yeast two-hybrid assay where INTS4 HEAT repeats are deleted at two HEAT repeat intervals. C. Same as panel B except regions at the C-terminus of INTS4 are being deleted. D. Same as panel C except smaller deletions from the C-terminus of INTS4 are tested. E. Same as panel D except mutations where five amino acids at a time are being changed to alanine in the context of the full-length protein. F. Western blot analysis of cell lysates transfected with the U7-GFP reporter after being treated with either control siRNA or INTS4 siRNA. In addition to reporter and siRNA, cells were also either transfected with empty vector (VA) or with RNAi-resistant INTS4 cDNA. G. Western blot analysis of cell lysates treated with control siRNA or INTS4 siRNA that were also transfected with the U7-GFP reporter and either empty vector or INTS4 cDNA that is RNAi-resistant. The INTS4 cDNA used were either wild-type or housing deletions in the HEAT repeats as labeled. H. Similar to panel G except alanine mutants of INTS4 were tested as described in panel E. Quantification is of 3 independent experiments and was quantified using a ChemiDoc scanner.

The results of the yeast two-hybrid analyses indicate that INTS4 associates with the INTS9/11 heterodimer but do not address the functional importance of this interaction. To address this, we utilized a reporter system that has been developed previously by our group called the U7-GFP reporter (Fig. 3F)(3,33,48,50). The U7-GFP reporter consists of the human U7snRNA gene upstream of a GFP ORF where GFP is only expressed upon transcriptional readthrough of the snRNA 3’-end processing site after INT knockdown. As an initial proof-of-principle, we knocked down INTS4 in human embryonic kidney HEK293T cells using two distinct siRNA targeting the INTS4 3’UTR and could detect ~75% knockdown efficiency at the protein level in both cases (Fig. 3F). While GFP expression from the U7-GFP reporter was undetectable in control siRNA-treated cells, we could detect significant accumulation of GFP after INTS4 knock down demonstrating that reduced expression of INTS4 leads to misprocessing of UsnRNA. We noted that the second INTS4 siRNA was slightly more potent to induce GFP expression and therefore, we used this siRNA for all further experiments (Fig. 3F). Importantly, we could restore GFP expression back to its original low levels in INTS4 knockdown cells if we re-expressed wild-type INTS4 mRNA containing the vector-derived 3’UTR rendering it RNAi-resistant. With these tools in hand, we proceeded to express cDNAs of INTS4 and measured their ability to rescue GFP expression of the U7-GFP reporter. Into cells treated with INTS4 siRNA, we expressed either full-length wild-type INTS4 or N-terminal truncations that removed HEAT repeats 1-2, 1-4, or 1-6. We observed that none of the HEAT repeat deletion mutants were capable of rescuing GFP expression and gave a level of GFP protein identical to that of cells treated with INTS4 siRNA and transfected with empty vector (Fig. 3G). Despite only ~50% reduction in interaction observed for INTS4 ΔH1-2 deletion mutant (Fig. 3B), this protein was not capable of any rescue in INTS4 depletion in cells suggesting that even a slight reduction in INTS4 association with INTS9/11 is sufficient to render the protein non-functional in cells. We extended this functional analysis to the C-terminus of INTS4 by expressing three alanine mutants: A/959-963 (mt1), A/954-958 (mt2), A/934-938 (mt5). The rationale is that mutant 1 and mutant 5 both retain association with INTS9/11 while mutant 2 has lost it. We observed that while mutants 1 and 5 rescue nearly to the same level as the wild-type protein, mutant 2 is significantly less capable of rescuing GFP expression levels (Fig. 3H). Therefore, we conclude that regions at the N-terminus and C-terminus of INTS4 that are required to associate with INTS9/11 are also required for INTS4 function in mediating 3’-end processing of pre-UsnRNA in cells.

### INTS4/9/11 heterotrimer constitutes the most critical subunits for snRNA 3’-end formation

Previously, it was demonstrated that Symplekin, CPSF73, and CPSF100 form a ‘core cleavage complex’ that is required for the 3’-end processing of both histone pre-mRNA and polyadenylated pre-mRNA (23,24). This model was based partly upon their physical association with each other but also upon the fact that RNAi-mediated depletion of these three factors, above all other members of the pre-mRNA processing machinery, led to the greatest amount of histone pre-mRNA misprocessing (24,31). Given that INTS4 binds to INTS9 and INTS11 once they heterodimerize, we posit that these three INT subunits form an analogous ‘core cleavage complex’ that is critical for the processing of UsnRNA. To attain functional evidence to support this model, we sought to knock down several members of INT and compare the subsequent impact on pre-UsnRNA 3’-end processing using our U7-GFP reporter. In this case, we chose to knockdown INT subunits in HeLa cells because knockdowns are typically more complete versus HEK293T cells. In addition to the two siRNA targeting INTS4, we also designed pairs of siRNA targeting: INTS1, INTS3, INTS7, INTS9, INTS10, INTS11, and INTS12. The rationale for choosing these other subunits to target is based upon experiments conducted in *Drosophila* S2 cells where we observed a wide range of UsnRNA misprocessing upon knockdown of the fly orthologues of these subunits (3). In S2 cells, depletion of INTS4/9/11 caused the most UsnRNA misprocessing, depletion of INTS1/7/12 caused moderate misprocessing, while depletion of INTS3/10 caused the least amount of misprocessing. In all cases, transfection of either siRNA into HeLa cells resulted in robust depletion of the targeted INT subunits (Fig. 4A). To assess the functional impact on pre-UsnRNA processing we transfected both the U7-GFP reporter and a plasmid encoding mCherry which contains a CMV promoter and SV40 polyadenylation signal to serve as transfection normalization to further increase the quantitative power of the U7-GFP reporter. As can be seen using fluorescence microscopy, the mCherry expression is the same in INTS4 siRNA transfected cells as in control treated cells but GFP expression is significantly higher upon knockdown of INTS4 (Fig. 4B upper panel). We conducted triplicate knockdowns using each siRNA and measured the amount of GFP expression using a fluorescence plate reader. As can be seen, depletion of INTS4, INTS9, or INTS11 in HeLa cells led to the highest level of GFP expression from the U7-GFP reporter while depletion of INTS10 and INTS12 led to only marginal GFP expression (Fig. 4B, lower blot and quantification). These results indicate that similar to the Symplekin, CPSF100, and CPSF73 core complex, depletion of each member of the INTS4/9/11 heterotrimer leads to the most robust amount of UsnRNA misprocessing when compared to depletion of several other INT subunits.

**Figure 4.**
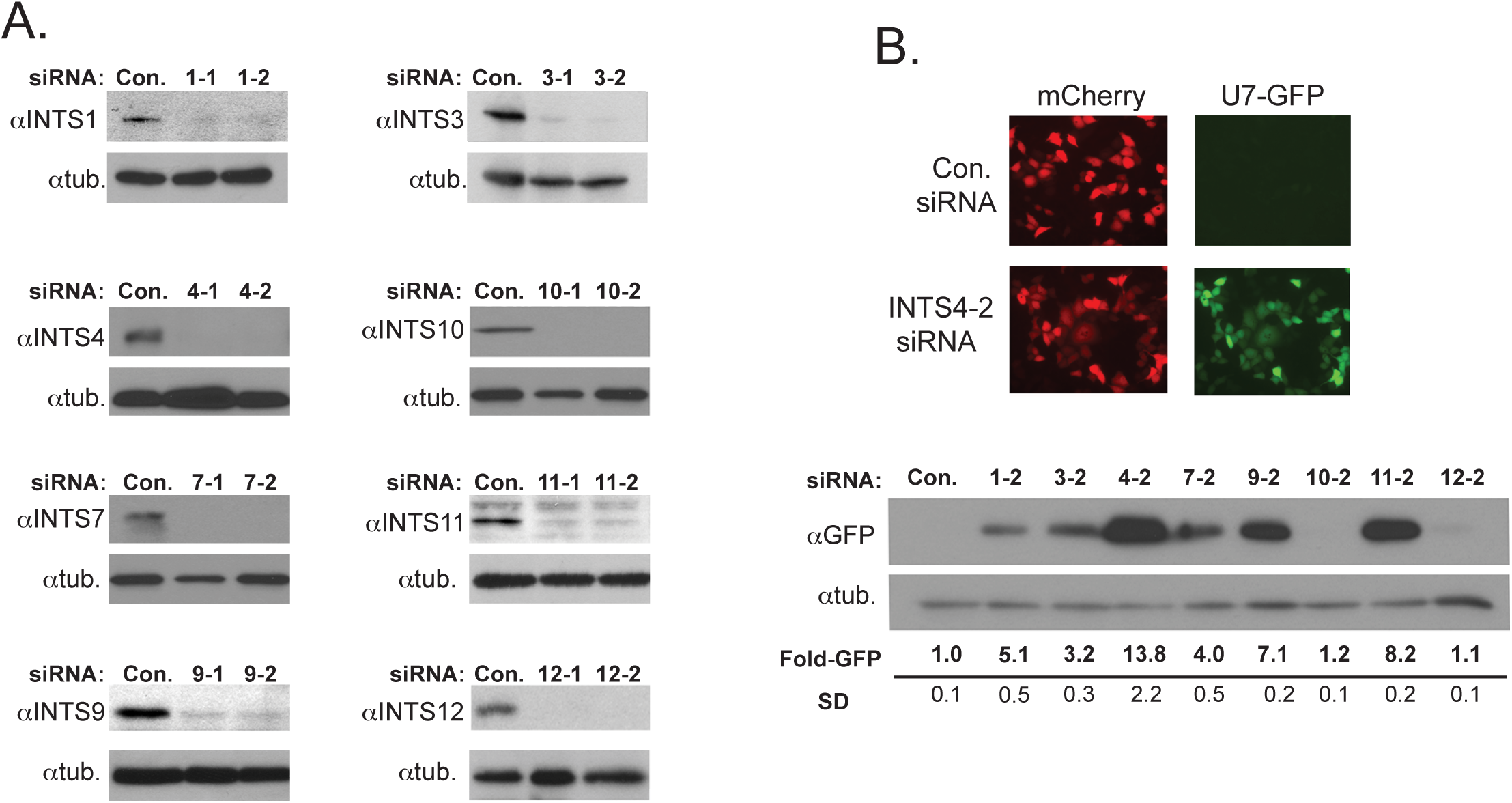
Depletion of INT subunits 4, 9, and 11 lead to the greatest amount of UsnRNA misprocessing. A. Western blot analysis of HeLa cells transfected with two different siRNA that each target eight individual INT subunits. Tubulin is used as a loading control. B. Upper panel-fluorescence microscopy image is representative of observations upon knockdown of INTS4. Lower panel is the results of fluorescence plate reader assays of triplicate transfections with either siRNA targeting INT subunits.

### The INT cleavage module is required for nuclear body integrity but is not a component of Cajal bodies

Previous work by Takata et al demonstrated that knockdown of INTS4 or INTS11 causes complete disassembly of Cajal bodies (CBs) and redistribution of the key structural CB proteins, coilin and SMN, to nucleoli (15). The conclusion drawn from this work was that INT is required for CB structural integrity, however, it is not known if INT subunits are physically present in CBs or if ongoing 3’-end processing of pre-UsnRNA precursors is the underlying requirement to maintain these nuclear structures (42). Using our current model that the subunits of the INT cleavage module are critical for INT-mediated pre-UsnRNA processing activity, we re-assessed the importance of INT in CB localization. As endogenous INT localization has not been evaluated in cells beyond INTS11, we first utilized a series of highly specific antibodies that were screened to determine where members of the INT cleavage module and several other subunits localize in HeLa cells (Fig. 5A). We observe that INT subunits 1 and 3 are strictly nuclear while INTS4, INTS7, INTS9, and INTS11 are both nuclear and cytoplasmic (Fig. 5B). Importantly, with the exception of some moderate overlap of INTS3 with CBs positively identified by CB marker protein coilin, we could never detect INT subunits colocalizing in CBs, however, in all cases we could observe INT localization in several prominent small foci adjacent to CB physical proximity.

**Figure 5.**
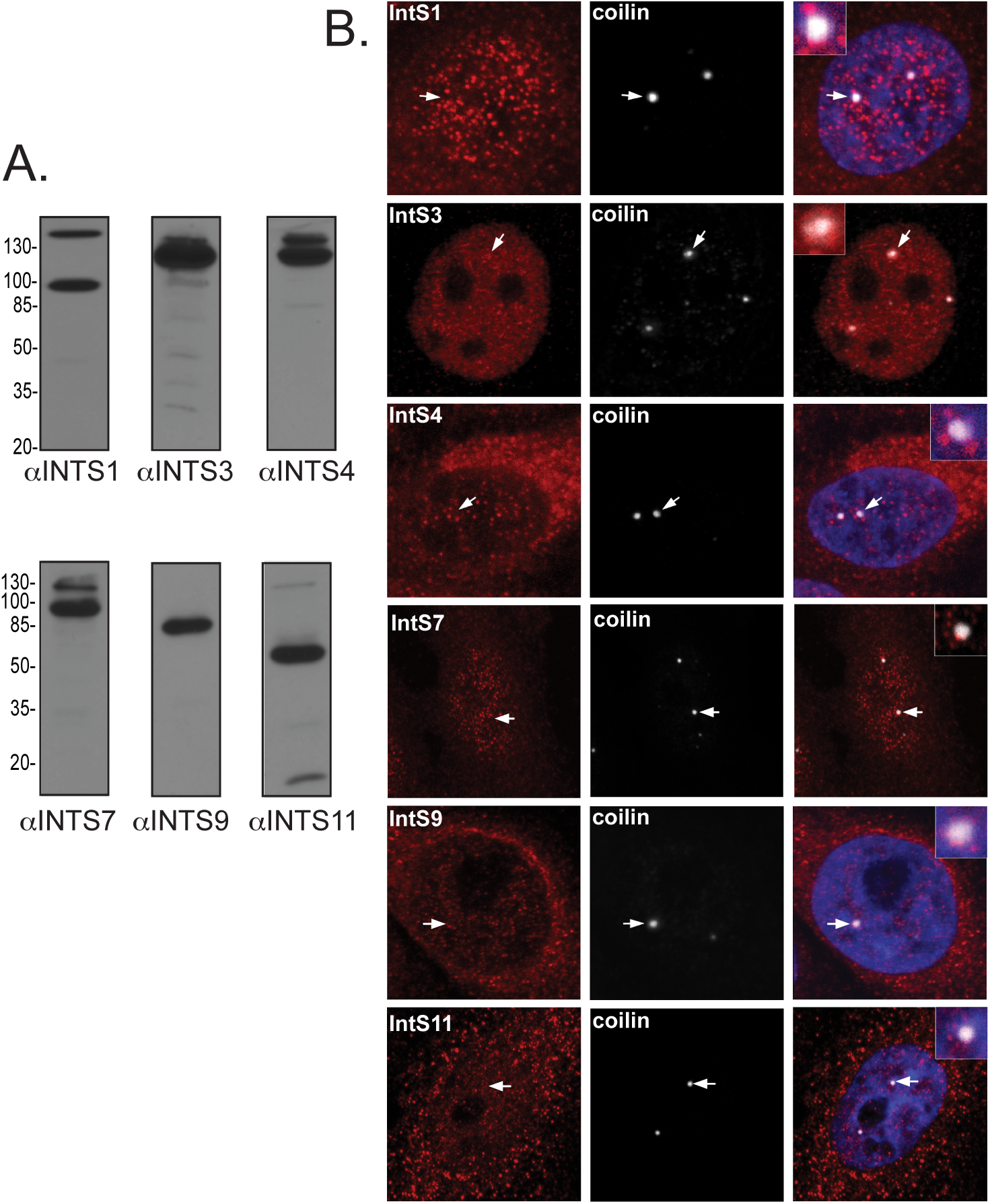
Localization of endogenous INT subunits. A. Western blot analysis of nuclear extract derived from HeLa cells. All bands on the Western blot have been confirmed using siRNA to be the labeled INT subunits. The lower band observed for INTS1 is a degradation product and not a nonspecific band. B. Confocal microscopy of HeLa cells fixed and immunostained with antibodies used in panel A and anti-coilin antibodies. DAPI (blue) is included in INTS1, 4, 9 and 11 to distinguish between nuclear and cytoplasmic INT subunit localization. Arrows indicate CBs surrounded by nearby IN subunit-positive foci, likely representing UsnRNA gene loci associated with CBs.

Given that none of the INT subunits that we tested provided evidence for robust colocalization within CBs, we hypothesized that INT proteins *per se* are not required for CB structural integrity but rather the ongoing production of processed UsnRNA transcripts through the cleavage activity of INT is required. Under this hypothesis, we predict the INT subunits that would be (most important to maintain CB structural integrity would be the same subunits that are the most critical for pre-UsnRNA 3’-end processing. Therefore, we repeated the RNAi experiments shown in Figure 5 and then fixed HeLa cells and subjected them to immunostaining for either coilin or Gemin2, a member of the SMN complex, to indicate the locations of CBs. CBs were readily detected in cells treated with either negative control siRNA or an siRNA targeting SLBP, which has been previously shown to have no effect on nuclear bodies (Fig. 6A)(51). Consistent with Takata et al, we observed complete disassembly of CBs and subsequent progressive relocalization into nucleoli after knockdown of INTS4 and INTS11 but surprisingly, the only other knockdown that caused a similar phenotype was of INTS9. We note that the coilin re-localization to nucleoli was confirmed by co-staining using antibodies raised against Fibrillarin, a 2’-O-methyltransferase enriched within the nucleolar dense fibrillar component (Fig 6B). We did not observe any significant change in CB integrity after knockdown of INTS3, INTS7, INTS10, or INTS12 and only a mild disruption of CBs after depletion of INTS7 (Fig 6C). These results demonstrate that only depletion of the Integrator cleavage module containing subunits (INTS4/9/11) causes complete CB structural disruption and indicates that the likely requirement for INT is to maintain CB integrity through INT’s function in continuous snRNA biogenesis. While the levels of CB protein components (e.g. coilin, SMN, Gemin2) were unchanged upon INTS11 knock down, we did observe significant co-depletion of both INTS9 and INTS4 further suggesting that these three subunits are tightly associated in cells (Fig. 6D).

**Figure 6.**
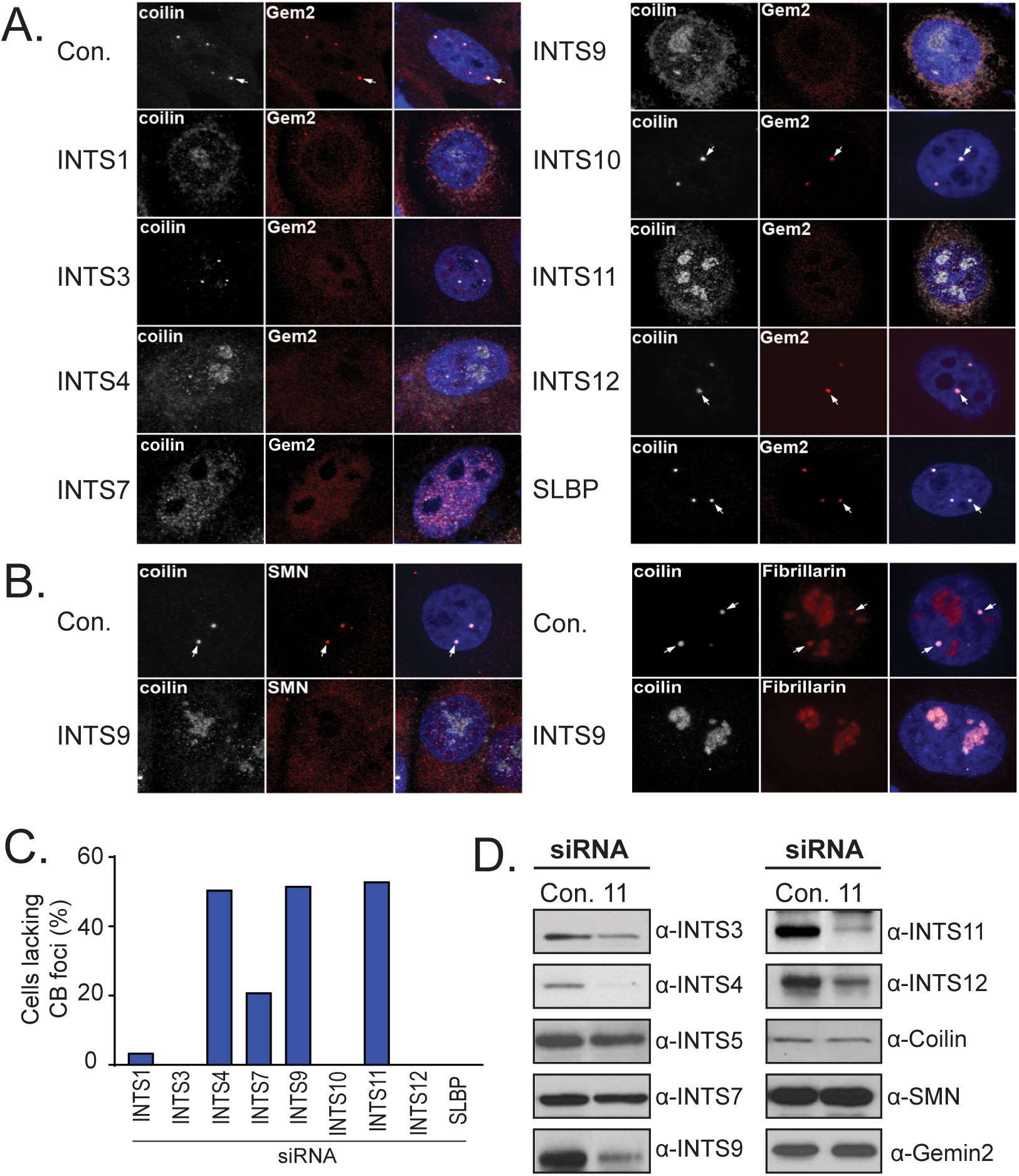
INTS4/9/11 are required for Cajal Body integrity. A. Confocal microscopy of HeLa cells fixed and immunostained with antibodies raised against either coilin or Gemin2, the member of the SMN complex, that have been transfected with siRNA targeting INT subunits or control siRNA (Con. or SLBP). Nuclei are visualized using DAPI staining (blue). Arrows indicate CBs in cells unaffected by specific INT subunit depletion. B. Similar to panel A except cells were only treated with either control siRNA or siRNA targeting INTS9 and stained for coilin, SMN, or fibrillarin. C. Quantification of 100 cells visualized from panel A demonstrating the percentage of cells with positive nucleolar coilin staining. D. Western blot analysis of cell lysates from cells treated with either control siRNA or INTS11 siRNA.

CBs are rich with a diverse range of small RNAs, most prominently spliceosomal snRNAs but also U7 snRNA, whose critical function is to mediate the 3’-end formation of histone premRNA during S-phase. This cellular process occurs in the HLB, which is formed on the two major human replication-dependent histone gene arrays (HIST1 and HIST2) and is frequently adjacent to CBs in human transformed cells. Previously, we have shown that CB integrity is required for histone gene expression (42) but the importance of INT to HLB formation and maintenance has never been explored. Given that depletion of INTS4/9/11 leads to the most significant amount of misprocessing of the U7-GFP gene reporter, we sought to determine what impact depletion of these subunits would have on these distinct nuclear bodies. HLBs can be detected through immunostaining with factors exclusively required for histone pre-mRNA transcription and processing (e.g. NPAT, FLASH, etc.) and can be observed to either localize associated with CBs or as individual nuclear bodies that are not associated with CBs. HLBs were readily detected in control or SLBP siRNA transfected HeLa cells and were visualized using antibodies raised against NPAT (Fig. 7A). The HeLa cells containing the greatest number of disassembled HLBs were those treated with siRNA targeting INTS4, INTS9, or INTS11 and to a lesser extent INTS1 (Fig. 7A/B). Complete HLB disassembly was not restricted to NPAT dispersion as HLBs visualized using FLASH were also lost upon knockdown of INTS9 (Fig. 7C). These results are strikingly similar to those obtained after immunostaining for coilin and suggest that disruption of both continuous spliceosomal snRNA and ongoing U7 snRNA 3’-end formation through depletion of the members of the INT cleavage module is sufficient to disrupt nuclear structures that are dependent upon these RNA for their proper function.

**Figure 7.**
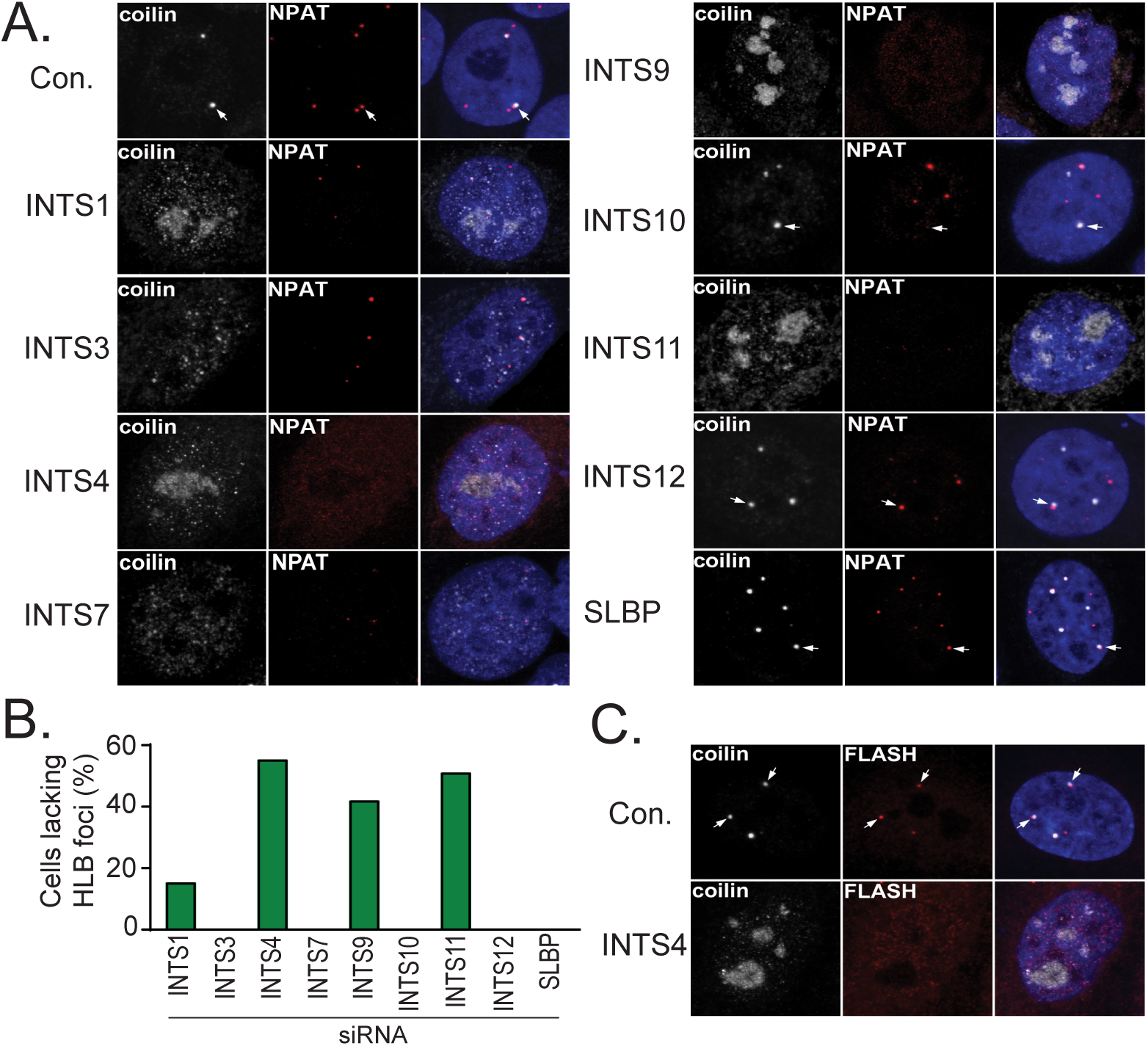
INTS4/9/11 are required to maintain histone locus bodies. A. Confocal microscopy of HeLa cells fixed and immunostained with antibodies raised against either coilin or NPAT, a specific histone transcription factor, that have been transfected with siRNA targeting INT subunits or control siRNA (Con. or SLBP). Nuclei are visualized using DAPI staining (purple). B. Quantification of 100 cells visualized from panel A demonstrating the percentage of cells lacking HLBs detected by NPAT staining. C. Similar to panel A except HeLa cells were only treated with either control siRNA or siRNA targeting INTS4 and stained for coilin and FLASH, a specific histone pre-mRNA processing factor, indicating that HLBs were truly disassembled in INTS4 kd cells.

## Discussion

In this study, we have identified INTS4 as the Integrator subunit that interacts with the INTS9/11 heterodimer. We believe that the evidence provided here supports a model where the INTS4/9/11 heterotrimer is the likely INT ‘cleavage module’ responsible for 3’-end formation of target RNA including both UsnRNA and eRNA (Fig. 8). While INTS11 contains the active site responsible for catalysis, its association with both INTS9 and INTS4 is critical to ensure proper UsnRNA biosynthesis. In turn, the integrity of the INT cleavage module helps to maintain the CB and HLB, which are two prominent structures that regulate essential cellular processes. These results uncover additional parallels between the cleavage and polyadenylation machinery and Integrator suggesting that the heterodimerization and scaffold association are important mechanistic components of both complexes to ensure accurate RNA cleavage.

**Figure 8.**
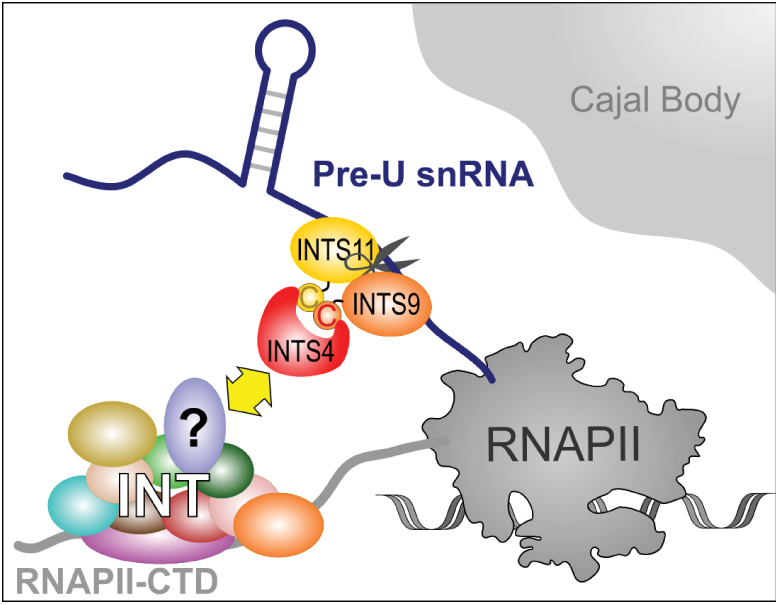
Model of the INTS4/9/11 Cleavage Module in UsnRNA 3’-end processing. Integrator associates with the CTD tail or RNA polymerase II to promote cotranscriptional processing of pre-UsnRNA immediately adjacent to the Cajal body. The INTS4 subunit utilizes both N- and C-terminal domains to interact with the INTS9/11 heterodimer to cause the cleavage of pre-UsnRNA. The heterotrimer of INTS4/9/11 associates with the rest of INT through interaction with a yet-to-be identified INT subunit.

The initial indication for a Symplekin function in pre-mRNA processing was derived from far western blot analysis using CstF64 as a bait protein (28), which was unexpected given earlier observations for Symplekin as a component of tight junctions (52). Despite these two opposing results, Symplekin can be purified from the nucleus associated with CPSF160, CPSF100, and CPSF73 (53). Symplekin can also be found in the cytoplasm associated with the same factors as well as CPEB and Gld-2 and was shown to be important to mediate cytoplasmic polyadenylation (54). These results indicate that Symplekin is a stable and essential component of the cleavage and polyadenylation machinery and, in particular, likely performs a function of scaffolding key subunits. Supporting this model, the *Saccharomyces cerevisiae* orthologue of Symplekin, Pta1, also is involved in cleavage and polyadenylation and performs an analogous scaffolding function through its association with members of both CPF and CF(30). Surprisingly, Symplekin was also found to be functionally required for the 3’-end formation of the replication-dependent histone mRNA (24,25,31,55). In the case of histone premRNA processing, Symplekin is thought to associate with CPSF73 and CPSF100 to form the Core Cleavage Complex (23). The CCC therefore incorporates into two distinct processing complexes involved in the cleavage of pre-mRNA that is subsequently polyadenylated or the replication-dependent histone pre-mRNA. While these two complexes share several components, they also have unique factors (e.g. SLBP, U7sRNP, PAP, etc.) suggesting that the CCC may associate differentially in order to cleave its two pre-mRNA substrates. Given that INTS9 and INTS11 are paralogues of CPSF100 and CPSF73, it is possible that INT has an equivalent CCC but, until now, there have been no reports to indicate which of the other INT subunits is performing the function of Symplekin.

Early indications of an important function for INTS4 were provided by studies conducted in *Drosophila* S2 cells. Previously, we depleted 12 of the fly INT subunits and measured the amount of GFP expression from the fly version of the U7-GFP reporter (3) and found strikingly similar observations to those presented here in human cells, in that, knock down of dINTS4 caused the greatest production of misprocessed UsnRNA followed by the levels measured after knock down of dINTS9 and dNTS11 (as well as dINTS1). Moreover, our structural analysis of the INTS9/11 heterodimer demonstrated that when these two proteins associate, there is a well-defined protein domain that is formed that has the strong potential to interact with other proteins given its β-saddle topology (34). The results of our modified yeast two-hybrid show that INTS4 will bind with INTS9/11 once they are associated but do not necessarily prove that INTS4 binds to the heterodimeric interface. As of yet, we cannot distinguish between the possibilities of INTS4 binding to the heterodimer domain or that INTS4 binds to a region of INTS9 or INTS11 that is only presented upon heterodimerization. Regardless, these data support the hypothesis that INTS4 is a scaffolding protein similar to Symplekin even though the two proteins are not paralogous based upon sequence comparison. A key difference between the two proteins is that the N-terminal HEAT repeats of INTS4 are somehow involved in binding to INTS9/11 whereas the analogous domain of Symplekin is dispensable to bind CPSF73/100. This difference could be related to the additional HEAT repeats that INTS4 appears to have relative to Symplekin and could suggest that INTS4 adopts a slightly different structure when it associates with INTS9/11. Indeed one of the canonical HEAT repeat containing proteins, PR65 (15 HEAT repeats) has been shown to adopt a half moon shaped structure where it can utilize its N and C-termini to associate with both the catalytic and regulatory subunits of protein phosphatase 2A in order to bring them in proximity (56). INTS4 could form a similar structure to PR65 or it could be that the HEAT repeats are simply required to adopt an overall fold that is necessary for its C-terminus to associate with INTS9/11. While Symplekin is clearly a scaffold for CPSF73/100, it also associates with other members of the cleavage and polyadenylation machinery (26,29) likely to promote targeting of the CCC to the site of pre-mRNA processing. By analogy, INTS4 is then the candidate to mediate incorporation of INTS9/11 to the remainder of the Integrator complex, but with which other INT subunit(s) is an open question (Fig 8). There are no INT subunits that obviously resemble members of the CstF complex or Ssu72, which are known Symplekin binding proteins suggesting that the incorporation of the INTS4/9/11 into INT is mediated by a novel interaction. Efforts are currently being focused to identify the INT subunit(s) responsible for recruiting the cleavage module into the complex.

Although a requirement for INTS4 and INTS11 for CB integrity has been established, it was not made clear if this requirement extends to all members of INT complex, if this requirement is derived from INT being a physical component of the CB, or if this is an indirect requirement of INT’s activity within the nucleus (i.e. extended pre-snRNA processing). The cellular localization of INT subunits has so far been restricted to either INTS9 or INTS11 or through the use of subunits expressed as fusions to GFP (10). Here, we assessed the endogenous localization of six INT subunits using a collection of highly specific antibodies and found that none of them were significantly localized within the CB. This indicates that the previous requirement of INT to maintain coilin accumulation in CBs is more than likely to be a downstream consequence of INT activity in the processing of pre-UsnRNA. A lack of a CB localization for INT is unexpected given the numerous reports of snRNA genes co-localizing in CB close physical proximity (37,42,57) and the fact that INT is thought to mediate UsnRNA 3’-end processing co-transcriptionally. These observations demonstrate that processing of pre-UsnRNA precursors by the INT cleavage module occurs adjacent to CBs rather than within them and that although INT subunits can be localized to UsnRNA gene bodies using chromatin immunoprecipitation (5), the association is likely to be transient or indirect. Furthermore, our data suggesting that the HLB is dependent on CB integrity for sustained function is noteworthy. We hypothesize that this arises from a direct supply of U7 snRNP from the CB to the HLB and this supply is vital to maintain HLB integrity. Altogether, our observations underscore the critical nature of the INT cleavage module to UsnRNA 3’-end formation in the context of nuclear structure maintenance and ultimately gene expression regulation.

## ACKNOWLEDGEMENTS

We thank Anupama Sataluri for her early work on this project and other members of the Wagner laboratory for helpful discussions.

